# Subthreshold violations of trajectory predictions are sensitive to TMS of Cerebellum CRUS I/II

**DOI:** 10.1101/2025.09.26.678469

**Authors:** Ellen Joos, Camille Scherer, Philippe Isope, Jack Foucher, Anne Giersch

## Abstract

Temporal prediction can help to follow a trajectory. In case of an error, the prediction can be adjusted. However, processing the error and adjusting the prediction can take time. What happens immediately after a prediction error, and can the processing of the prediction be modulated? We use a newly found illusion based on moving squares and requiring trajectory regularity to be elicited. We examined the conscious consequences of a sub-threshold manipulation of the square trajectories, and transcranial magnetic stimulation (TMS) on the cerebellum (right CRUS I/II) to study the modulation of the processing of the trajectory manipulations. The TMS was a typical intermittent theta-burst stimulation, but only one sequence of around 3 minutes, compared with a placebo stimulation. The trajectory manipulation had a reliable effect on the illusion, even though the illusion emerged within less than 100 ms after the trajectory manipulation. The results suggest that the prediction is temporarily stopped after the trajectory change. The illusion was accompanied by EEG signals whose amplitude was modulated by TMS on the cerebellum, at least in those participants who received verum TMS after having performed the task three times. Those EEG signals resembled a late LPP (Late Positive Potential). As LPP spontaneously decreased over time, the results suggest the effect of TMS may represent a reinstation of the EEG consequences of the prediction error, i.e., a modulation of its significance.

## Introduction

Predicting future information in order to adjust one’s own behavior, e.g., when a car is heading towards us, can be crucial in everyday life. Current theories postulate that the brain detects regularities in the sensory input and forms predictions accordingly in an internal model (Wolpert et al., 1998). A difference between the model and the actual sensory information leads to so-called “prediction errors”, which signals the need to adjust behavior (Giersch et al., 2016; Shadmehr et al., 2010; Sokolov et al., 2017). However, what makes a prediction error pertinent or not is not always straightforward. In everyday life, it is frequently the case that there are unexpected glitches, even when events unfold as usual, but those glitches are sometimes best ignored. For example, it is unclear if subthreshold irregularities in a car trajectory in front of us are or not significant (subthreshold irregularities means irregularities that are consciously perceived in less than 50% of the cases). It has been shown many times that subthreshold prediction errors are processed, but their impact on perception and whether or not their processing is modulated is unclear. What happens if the prediction error has been processed in the brain, e.g., the irregularity in the car trajectory in front of us, but we are not aware of this irregularity? If the trajectory prediction is affected, what is the consequence on conscious perception during the interval between the prediction error detection and the adjustment? Is it possible for the response to prediction errors to be modulated? Here we explore (1) whether sub-threshold irregularities impact the prediction of a movement trajectory within 60 ms after the irregularity and its effect on conscious perception, and (2) whether the processing of irregularities can be modulated by stimulating the cerebellum, that has a growing literature showing its involvement in temporal prediction and millisecond-level processes (Sokolov et al., 2017).

### Modulation of time prediction errors

A number of studies have suggested that the processing of prediction errors can be modulated. The first factor is the regularity of sensory information itself (Barascud et al., 2016; Friston, 2005, 2009). A prediction error is processed as such only if information is regular enough. The precision of the detected prediction errors increases with this regularity. For example, larger neural correlates’ amplitude is observed in the context of high rather than low regularity (Garrido et al., 2013). Conversely, when the prediction error becomes frequent, it is learned and ignored (Brooks et al., 2015). In addition to regularity, attention also can increase the precision of the prediction errors (Feldman & Friston, 2010). However, the role of these factors has mainly been explored with detectable oddballs that lead to prediction errors emerging at the conscious level.

The consequences of supra-vs. subthreshold prediction errors may differ. It has been suggested that in case of small prediction errors, the internal model is updated, whereas in case of a large error, a new model is created. Oh and Schweighofer in humans (Oh & Schweighofer, 2019) and Spaeth et al. in mice (Spaeth et al., 2022) largely confirmed this hypothesis, but interestingly both results also showed interindividual differences. Moreover, in real life, it is rare that there is only one prediction error. Tabas & Kriegstein (Tabas & Kriegstein, 2024) added a level of complexity of prediction error processing by exploring the modulation of prediction error signals in case of multiple, contradictory signals. They showed that contrary to prior beliefs, the integration of multiple prediction errors is not linear. There might thus be multiple ways to modulate the processing of prediction errors.

### Sub-threshold prediction errors

We used sub-threshold irregularities because their processing was most likely modulated (independent of conscious realization and attention modulation). A second motivation for the focus on sub-threshold irregularities was the observation of abnormalities in the processing of asynchronies and delays in schizophrenia, that affected specifically the millisecond, sub-threshold time manipulations (Foerster et al., 2021; Lalanne et al., 2012; Marques-Carneiro et al., 2021). These results suggest that there might be some inter-individual differences at this time level specifically, i.e., at the level of milliseconds. An abnormal sensitivity to low-level prediction errors has also been proposed in autism (Van de Cruys et al., 2014). According to the HIPPEA model, it might be related to a failure in the regulation of the prediction error precision. We thus have several reasons to explore the impact of small irregularities and the modulation of their processing. In the present study, we explore this question in neurotypicals, but the results are intended to shed light on mechanisms that may have a pathophysiological role in schizophrenia and autism.

### The role of cerebellum

In this context, the cerebellum is an obvious target (Giersch et al., 2016; Parker, 2016; Picard et al., 2008), and this for several reasons. First, even though the cerebellum is mainly known for its role in motor functions and in motor predictions (Paulin, 1993; Person, 2019), its involvement in sensory and cognitive mechanisms is now largely recognized (Andreasen et al., 1999; Argyropoulos et al., 2020; Leiner et al., 1993), especially its role in the general formation of predictions (Popa & Ebner, 2019; Sokolov et al., 2017). Moreover, the cerebellum likely plays a role in the definition of the prediction errors precision, even though it is not the only area involved (Alexander & Brown, 2019; Man et al., 2024). The cerebellum has been shown to code prediction errors, like e.g., unexpected self-motion (Brooks & Cullen, 2013). Most importantly, it is also known to be involved in time processing at the level of milliseconds (Coull et al., 2011; Ivry & Spencer, 2004), which makes it a choice target when considering sub-threshold trajectory manipulations. It should be noted also that the cerebellum is hypothesized to play a role in those pathologies showing sensitivity to low-level prediction errors, such as schizophrenia (Andreasen, 1999; Cattarinussi et al., 2024; Feng et al., 2024; Jensen et al., 2024; Kang et al., 2024; Laidi et al., 2019; Mehta et al., 2024), or autism (Goodwill et al., 2023; van der Heijden et al., 2021). Regarding our specific aim, i.e., trajectory manipulation, the literature also suggests the involvement of the cerebellum. Several studies have explored the impact of TMS targeting the cerebellum (Mioni et al., 2020), although mainly on duration perception. Most importantly, several studies suggest a role of the cerebellum in the processing of moving object trajectories. For example, cerebellar patients are impaired at discriminating the velocity of moving objects (Ivry* & Diener, 1991) or at discriminating motion from noise (Thier et al., 1999). Although there are visual cortex areas specialized in motion processing, like MT and MST, predicting future trajectories of external objects to avoid collisions is of paramount importance. It is thus not surprising that the cerebellum, and especially its lateral part (Crus I, Lobule VII) is involved in the detection of motion and the prediction of temporo-spatial information in the case of visual trajectories, both in humans (Baumann et al., 2015; Baumann & Mattingley, 2010; O’Reilly et al., 2008) and cats (Cerminara et al., 2009). For example, based on the current state of the body and motor commands originating in the cerebral cortex, cerebellar internal models are updated through the modification of synaptic transmission between granule cells and Purkinje cells (Cullen, 2023; Spaeth et al., 2022; Streng et al., 2022; Wolpert et al., 1998). This plasticity is controlled by the climbing fiber pathway as it conveys error and/or reward signals to the Purkinje cell (Hull, 2020). Error predictions can therefore be encoded by Purkinje cell discharge, which is transmitted to the cerebral cortex via the cerebellar nuclei and the thalamus (Li & Mrsic-Flogel, 2020; Ramnani, 2006). The literature thus provides grounds to explore the modulation of sub-threshold prediction errors, beyond attention and statistical regularities manipulations. This justifies to target the cerebellum with TMS when the task involves trajectory predictions.

### Rationale for the task used to investigate adjustments to sub-threshold trajectory manipulations

We used a new illusion discovered by Jovanovic et al. (Jovanovic et al., 2023). The task involved visual stimulus trajectories, which were manipulated at the subliminal level. Furthermore, we evaluated the impact of both the trajectory manipulation and of a TMS stimulation targeting the right cerebellum on subjective perception and on EEG signals.

On each trial, two squares move towards each other on a screen. In case of a long contact duration of the two squares, participants easily identify that the squares touch (Figure 2). In case of short contact durations (17ms and 33ms), however, the contact is not perceived, but rather an illusory gap. The illusion does not result from the brevity of the contact, since the contact is visible in several conditions even for 17 ms contacts, e.g., when squares are empty rather than filled in. Several control experiments led to the interpretation that the illusion originates from the violation of the low-level edge contrast predictions. In detail, two black squares move towards each other on a gray background. The high-level, cognitive prediction would be that the squares touch in the middle of the screen, which is not what participants perceive in case of very short contact durations. It is rather the low-level prediction that seems to evoke the illusory gap through the regularity of the continuously moving edge contrast shift along the trajectory. According to this low-level prediction for position x+1, the inner edge of the left square should evoke a prediction of black on the left and gray on the right side of the edge, whereas for the right square it is gray on the left and black on the right. Even though the two squares touch, gray is predicted at the center of the screen. This illusory gap is what participants perceive, suggesting that the low-level predictions lead to this subjective perception.

As already emphasized, the regularity of prior information is important for a prediction error to emerge. The question is how a perturbation of this regularity affects not only the prediction but also the illusion. When the perturbation is sub-threshold and undetectable consciously, it should be possible to ignore it. If not, the prediction should be affected. For example, if the squares accelerate, the prediction of the next squares’ position should be adjusted. However, adjusting the prediction should take time, and since the contact occurs in less than 100ms after the trajectory perturbation, we can expect to see the result of a lack of prediction rather than an adjusted prediction. Preliminary experiments confirmed an impact of an acceleration, when occurring around 30 milliseconds before the end of the trajectory (when the perturbation occurred earlier, the trajectory was regular again and there was no impact). If replicated, it would be an argument for the link between the prediction and the illusion, and would illustrate the perceptual consequences of prediction errors. Given the role of the cerebellum in the processing of prediction errors at the level of milliseconds, and in the processing of trajectories, we further tested our hypothesis with non-invasive brain stimulation. Transcranial magnetic stimulation (TMS) is a very effective and promising tool to alter neural processing. We targeted the CRUS I/II regions due to their role in trajectory prediction and perception, and their link with the fronto-parietal cortex (Guell & Schmahmann, 2020; King et al., 2019).

We measured electrophysiological (EEG) in addition to behavioral responses. Several EEG signals have been put in relation with the processing of prediction errors, like e.g., the N1 or MMN evoked potentials (Lieder et al., 2013; Pinheiro et al., 2019). Those signals have been observed in oddball paradigms, when the prediction is not only rare but emerges at the conscious level. It has been suggested that some degree of consciousness is required for a MMN to be observed (Flynn et al., 2017). Given the subconscious character of our manipulations, a MMN was not expected (but see (Liaukovich et al., 2022)). However, other signals may be observed. An update of predictions should follow a subliminal prediction error, and late signals have been described up to 1 second after a prediction error, like a decrease in beta oscillation, a feedback related negativity, or a late positive potential (Burnside et al., 2019; Chao et al., 2018; Hajcak & Foti, 2020). Given the new approach, it was difficult to predict what type of EEG results would be observed exactly, but EEG changes proved to be especially sensitive to the effect of TMS.

To summarize our objective and hypotheses, the present study documents the impact of sub-threshold, millisecond-level perturbations on conscious perception, and seeks to determine how it is modulated by collecting a behavioral measure, EEG correlates, and by stimulating the cerebellum by means of TMS. The modulation of the processing of prediction errors was explored first by verifying the stability of behavioral and EEG results, independent of TMS. It can be expected that the processing of sub-threshold errors may vary across time and training, and we tested the stability of behavioral and EEG results. Finally, we used TMS on the cerebellum to verify if the impact of the prediction errors on the illusion rate and on EEG correlates can be reinforced.

## Methods

### Participants

We recruited 24 healthy young adults (12 females, mean age ± SD: 24 ± 5 years) that were mainly students of the University of Strasbourg. After having received detailed explanations about the experiment, participants gave their informed written consent and the study was conducted in accordance with the ethical agreements of Helsinki (Association, 2000) and approved by the ethical committee CPP IV Est (N° IDRCB: 2016-A00106-45).

### Experimental procedure

Participants came to the lab 4 times (see Figure 1). Once written consent had been obtained, we checked for exclusion criteria: (history of) neurological or psychiatric diseases, drug abuse, usage of psycho-active substances, take-in of benzodiazepines, cannabis consummation or other hallucinogen substances, decreased visual acuity (Bach, 1996), counter indications for MRI or TMS (presence of permanent ferromagnetic body, pacemaker, prosthesis, vascular clip or stent, etc.), pregnancy or breast-feeding. In the same meeting, we additionally ensured right-handedness via a handedness test (Oldfield, 1971) and measured neuropsychological abilities via the French National Adult Reading Test (Mackinnon & Mulligan, 2005), the CPT-AX (Cohen et al., 1999), a neurological test for soft signs in Schizophrenia (Krebs et al., 2000), which were within norms for all participants.

**Figure 1.**
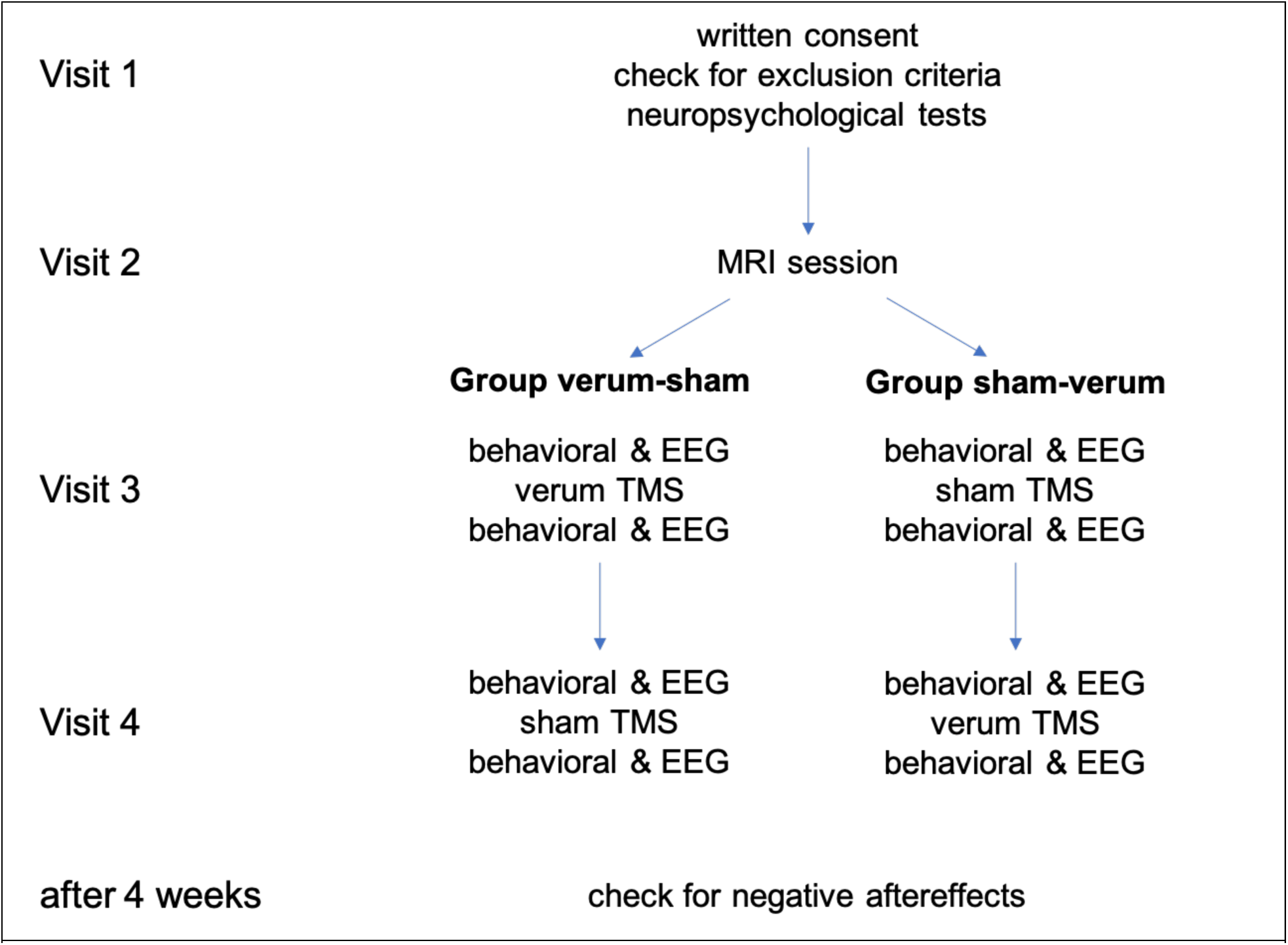
Experimental Procedure – Participants were invited to the lab 4 times: In visit 1, we explained the procedure, obtained the informed written consent, checked for exclusion criteria and performed neuropsychological tests. In visit 2, we measured participants in the MRI scanner, mainly to inform the TMS robot in the later intervention. We randomly assigned a group to each participant (counterbalanced between them), where in group verum-sham, participants received verum TMS on visit 3 and sham TMS on visit 4. In group sham-verum, participants received sham TMS on visit 3 and verum TMS on visit 4. Before and after the interventions, we measured behavioral and EEG responses. 4 weeks after concluding the experiments, we called participants to ensure that they did not experience any negative aftereffects.

In the second meeting, participants were invited to a session in the MRI machine (Siemens Magnetom Verio 3T with a 32 channels head coil). We obtained anatomical data in order to later inform the neuronavigational system of the TMS robot. During this MRI session we also obtained functional MRI of the collision task and of another temporal prediction task (data not shown here).

In the third and fourth session, we measured behavioral and electrophysiological (Biosemi EEG system with 64 channels) data before and after transcranial magnetic stimulation (TMS) intervention. It should be noted that participants additionally performed a short version of the illusion task immediately before and after the intervention. Although this yielded no significant result, it may have facilitated the effect of the TMS on the responses observed in the illusion task (Silvanto et al., 2007). The intervention was either the verum transcranial magnetic stimulation (TMS) or sham TMS (see more details below). Verum and sham TMS were performed on separate meetings and their sequence was counterbalanced between participants. We ensured a minimum of 6 days between the two meetings in order to prevent spill-over effects of the verum TMS on the sham TMS. The employees at the TMS platform who performed the intervention were necessarily aware of the type of intervention, but the experimenters were not and they were also not present in the room during the intervention. It is to be noted that no participant had any nausea or muscle contracture following the TMS targeting the cerebellum. The experimental part (before and after intervention) took place in the psychiatric hospital of the University of Strasbourg, while the intervention was performed at the CEMNIS, Strasbourg. The two locations are 200m apart from each other and the experimenters walked with the participant from one location to the other. Several EEG electrodes had to be unplugged in order to enable the TMS intervention, but the rest of the EEG setup remained on the participants’ head during the whole meeting.

As a final step, we called participants 4 weeks after completing the experiment and none of the participants showed any sign of (negative) after effects.

### Verum TMS and sham TMS

The transcranial magnet stimulation was a single intermittent theta-burst stimulation (iTBS) with 10 bursts of 3 biphasic pulses at 50 Hz, repeated at 5Hz and performed 20 times with 8s burst intervals (Huang et al., 2005). The total duration of the stimulation was 3min 22seconds. Stimulations were executed via the Axilum Robotics TMS robot with the neuronavigation system Localite that was informed by the individual anatomical data obtained during meeting 2 (at least 7 days before meeting 3). We used MagVenture as the stimulator and a figure of 8 coil.

Stimulating the Cerebellum with TMS is difficult due to the depth of the structure. We aimed at stimulating the border between regions Crus I and Crus II on the right side. We selected the right side because prior studies showed larger effects on sub-second perception duration or sensorimotor integration in the case of a stimulation on the right than on the left cerebellum hemisphere (Bijsterbosch et al., 2011).

In order to account for individual scalp properties, we selected two possible targets, while aiming to stimulate the most lateral and more anterior part of the border (target 1). If this was not possible, a location more lateral and closer to intermediate region was stimulated (target 2). Target 1 was stimulated in 19 participants, while target 2 was stimulated in 5 participants. Target location did not have an effect on behavioral or electrophysiological measures and in the following results are thus presented across both targets (see Supplementary Material S1).

We did not use the active motor threshold to determine the stimulation intensity, because the motor cortex is functionally and spatially very far from the Cerebellum. Instead, the target location was used to determine the stimulation intensity. Participants were instructed to tap their fingers from the small finger to the thumb sequentially and in a smooth manner and to repeat this movement constantly. Single pulses were applied to Crus I/II and intensity was continuously increased until the movement was interrupted by the pulses, which we label the cerebellar threshold. For the iTBS intervention, the stimulation intensity was 100% of this cerebellar threshold. According to these parameters, the intensity used for our group should have been 59.3 ±11.5% (mean ± SD) of the stimulator’s maximum output. However, six participants experienced tolerance issues (muscle pain) during the verum stimaution and they required a reduction in intensity (10.3 ± 10.1%), resulting in an actual intensity of 56.8 ± 10.7% of the stimulator’s maximum output.

In the sham TMS session, the coil was reversed such that the two currents cancel each other out. Participants did not know whether they received the verum or the sham TMS intervention. We could, however, not prevent that participants might have felt a difference due to (neck) muscle activation.

### Equipment and apparatus

Participants were seated in a dimly lit and quiet room. A chinrest ensured the distance between head and screen of 67cm. The stimuli and the procedure were programmed using MATLAB (version 2009b) on a HP Compaq 8100 Elite computer and presented on a Sony CPD-520GST Trinitron monitor with a refresh rate of 60 Hz and a resolution of 1280×1024 pixels. Manual responses were recorded using the keys “f” and “j” on a standard keyboard.

### Stimuli and experimental design

We presented the recently developed collision task (Jovanovic et al., 2023) as illustrated in Figure 2. In this visual illusion two black squares (1×1°VA, 0.07 cd/m²) were presented on a light gray background (8.9 cd/m²). They appeared 1° visual angle left and right of the middle of the screen. In the following 500 milliseconds, the squares moved towards each other until their inner edges touched at the middle of the screen. The squares would then stay on the screen and disappear 17, 33, or 200 ms after their contact. After a 250ms waiting period, a response time window started in which participants were instructed to press one key in case that the squares touched and another key in case that the squares did not touch. The response time window was adapted to the contact duration, i.e., 1733 ms response window with 17 ms contact duration, 1717 ms response window with 33 ms contact duration, and 1550 ms response time window with 200 ms (no) contact duration. Afterwards, a fixed inter-trial interval (ITI) of 500ms was presented. The total duration of one trial was always 3000ms.

**Figure 2.**
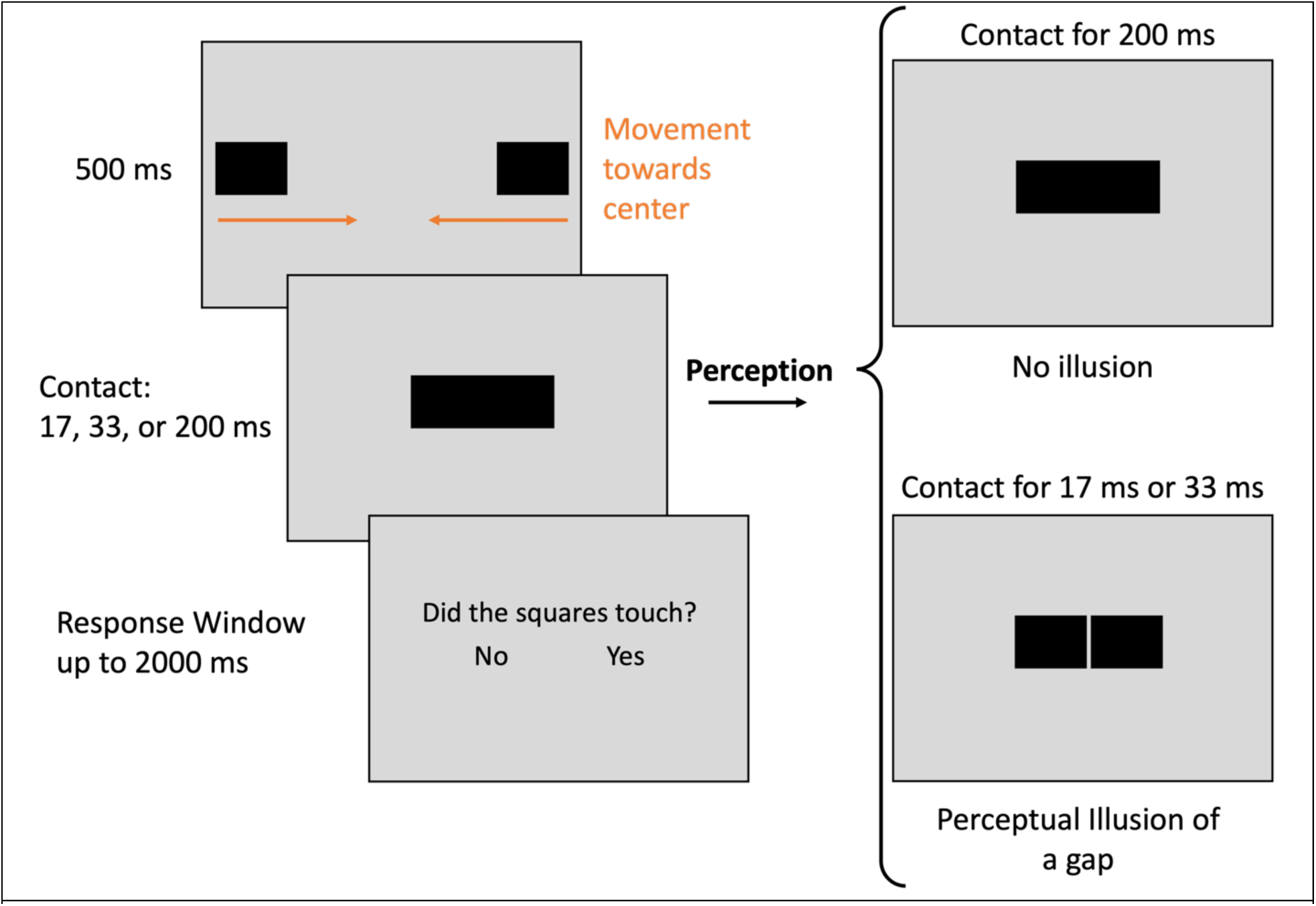
Two squares appear on the screen. They move towards the center until the inner edges of the squares touch for variable amounts of time (17, 33, or 200ms contact duration). The 200ms long contact duration was clearly visible and was used to ascertain that the participant followed instructions. With short contact durations (17 and 33ms), the touch of the squares is not necessarily perceived, which results in the perception of an illusory gap. Participants were asked to report whether they perceived the squares touching or not in a response time window following the contact.

Importantly, the 33ms long contact of the squares is not necessarily perceived by the participants and thus results in the illusory perception of a gap between the two squares. Due to a previous study (Jovanovic et al., 2023), we expected an illusion perception rate of roughly 60% in the 33ms trials (see Supplementary Material S2). Like in preliminary studies, we privileged 33ms trials (68% of all 300 trials), because it allowed to reliably measure both an increase and a decrease in illusion perception rate, i.e., any modulation of the illusion rate. The other 32% of the trials were evenly split between a contact duration of 200ms and a contact duration of 17ms. The 200ms trials served as control conditions, because with this long contact duration no illusion is perceived. In order to check that participants understood both key-meaning associations, we presented half of the 200ms trials with the squares touching in the middle of the screen, which is easily detectable as a contact. The other half of the 200ms trials was presented with the squares stopping their trajectory one frame, i.e., 1 pixel, before they would touch, which is easily detectable as a gap. The 17ms trials were presented because previously, an illusion perception rate of roughly 90% was measured with the 17ms trials. Those trials thus served as a control to ensure that participants understood the task in a condition in which the illusion was perceived with a high probability (as compared to the 200ms trials).

In the trials in which the trajectory was standard without perturbation, the squares moved 30 pixels within 500ms, resulting in 1 pixel per frame. Trajectory manipulation was introduced in some trials to disturb low-level predictions. In case of a perturbation, the squares moved 2 instead of 1 pixel within 1 frame. To avoid ending the trajectory on this 2-pixels-jump, the square moved one pixel more, meaning that the perturbation occurred 2 frames (33ms) before the contact. The squares then stayed on the screen for 17, 33 or 200 ms, and then disappeared. In most of the cases, this perturbation was not consciously perceived by the participants, as checked in the control condition presented in the supplementary material S3 (Fig. S2 b)).

In the first block, we showed 100 trials that only contained non-perturbed trials (68 33ms-trials, 16 17ms-trials, 8 200ms-with contact, 8 200ms-without contact). In the second and third block, we presented 100 trials each. In each of these blocks, half of the trials containing non-perturbed and the other half of the trials containing perturbed trials.

The sequence of trials was determined using OptSeq2 (Dale, 1999) (10000 randomizations, optimal result taken for all participants; software available at https://surfer.nmr.mgh.harvard.edu/optseq/).

### Behavioral analysis

We calculated the illusion perception rate, i.e., the percentage of reports that the squares did not touch (which equals to an illusory gap). Two participants had less than 75% correct responses in either of the 200ms control conditions and were thus excluded from further analyses. Further, three participants had to be excluded due to abnormal perception in one of the trial types, i.e., one participant had 20% illusion perception rate instead of roughly 90% in the other trial types, and another 2 participants showed a floor effect of the illusion (less than 20% illusion perception rate) in the non-perturbed trials of the mixed blocks. The possibility to respond was given 250ms after the offset of the squares and thus we did not have any constrains for reaction time values.

For the behavioral analyses, we analyzed only the trials with 33ms contact duration at trial N. The experimental design included different types of trials (non-perturbed in two different blocks and perturbed trials) and several time points of measurement, i.e., before and after either verum or sham TMS interventions. Further, our design enabled a within comparison of the factors *Intervention* (verum or sham TMS) and *Time Point* (before or after the intervention) effects. As a result, half of the participants received the verum TMS in their first intervention meeting and the other half received sham TMS in their first intervention meeting. To investigate a possible influence of order of the sessions on the results, we additionally tested for the factor *Session Order* (Session 1 verum TMS, i.e., verum-sham, or Session 1 sham TMS, i.e., sham-verum). To facilitate the readers understanding of our analyses, we describe the details in the respective results part.

We used univariate Bayesian analyses with non-informative priors to investigate the above-mentioned effects (see also (Arrouet et al., 2022)). All data was non-symmetrically distributed and thus was fitted with a beta regression. The probability for a difference of the respective comparison is denoted by e.g. *Pr*(verum TMS > sham TMS), i.e. that the performance is better with the verum TMS than with the sham TMS. For interactions, the probabilities are written as follows: *Pr*(OR > 1) with OR exp*^B^* being the coefficient of the interaction in the beta regression. In case of meaningful interaction effects, we calculated pairwise comparison to isolate the origin of the effect. We define meaningful effects with values that are *Pr* > 0.975 or *Pr* < 0.025 since both are equivalent, as *Pr* (A > B) = 1 − *Pr* (A < B). All Bayesian analyses were conducted using the R packages jags (Plummer, 2003) and brms (Bürkner, 2020).

### Electrophysiological analysis

#### EEG recordings and pre-processing

We measured the EEG with the ActiveTwo 64 channel BioSemi system with active silver/silver chloride electrodes. EEG data were digitized with a sampling rate of 2048 Hz. The data was offline down sampled to 512 Hz and digitally filtered with a low-pass at 25Hz and re-referenced to the common average. Data analysis was executed in Python using mne (Gramfort et al., 2013; Larson et al., 2024).

We detected bad electrodes by manually selecting those electrodes that showed an abnormal power spectral density. Blinks were detected using the mne ICA implementation. To do so, we filtered the data from 1-30 Hz, broke it down into epochs of 1s, and identified components as representing blinks when their correlation with electrodes Fp1 and Fp2 were equal or higher than a correlation coefficient of 0.8. In 14 out of 96 cases (24 participants, 4 EEG sessions each, i.e., before and after the two types of intervention) we had to manually correct the identified blink components. After removing those blink artefacts from the individual EEG data, we excluded trials from analysis when reaching an artefact threshold of ± 150 *μ* V. Amplitudes were measured relative to baseline, which was defined from 100ms before until the onset of the square’s trajectory. Trials were calculated until 2900ms after the trajectory, i.e., until the end of the ITI. EEG data was sorted the same as in the behavioral analyses. We set a minimum of 20 trials per condition, participant, and trial type and averaged across illusion and no illusion trials. Due to problems during data acquisition, we had to exclude three out of the originally 24 participants from the EEG analyses.

#### ERP analyses

The data were separately analyzed and averaged for each participant and for each EEG electrode using the onset of the trajectory as a time reference and then averaged across participants to obtain the grand mean data.

This study is somewhat explorative, since we had never measured EEG when presenting this visual illusion. We visually inspected the whole range of grand mean ERP data and found several temporal and spatial regions of interest. We selected the spatial and temporal region of interest that yielded the most meaningful results in terms of intervention influence, i.e., a difference between pre vs. post compared between verum TMS and sham TMS. This exploratory analysis revealed a differential effect of intervention at electrode CP4 in the time range from 0.15s until 2s after the contact of the squares, i.e., the whole response time window. We conducted the same statistical comparisons as described in the behavioral analyses and the results section for these ERP amplitudes at CP4.

#### Correlation of behavioral and ERP results

We calculated the correlations between behavioral and ERP results using Spearman rank correlations in R. This was done separately for the different conditions (non-perturbed trials in the first block, in the second/third block, and perturbed trials in the second/third block), types (verum TMS and sham TMS), and time points (pre vs. post) of interventions. Participants excluded from the behavioral analyses were not the same as those excluded in the EEG analyses. Reliable data in both measures was found in 16 participants, which were then used for calculating the correlations. We corrected for multiple testing using the Bonferroni method.

## Results

### Trajectory perturbations and group influence responses to the illusion

Our first aim was to verify whether the trajectory perturbation impacted the illusion, irrespective of the TMS interventions. To do so, we considered only data from the morning sessions. We investigated behavioral and EEG differences between trial types: 1) non-perturbed trials of the first block, in which solely non-perturbed trials were presented, 2) non-perturbed trials of the second and third block, i.e., those blocks in which non-perturbed and perturbed trials were mixed, and 3) perturbed trials of the second and third block (factor *Trial Type*). We analyzed the effects’ stability over time by comparing results from the morning of session 1 with that from the morning of session 2 (factor *Session*). However, there was either a verum or a sham TMS session in between the morning sessions, that might have influenced the data. For this reason, we took into account the order. With the same reasoning, we additionally tested for a possible difference between the two groups (factor *Session Order*), i.e., group receiving verum TMS on session 1 (group verum-sham) or on session 2 (group sham-verum).

We found that the illusion perception rate was clearly higher for the non-perturbed trials than for the perturbed trials [OR = 0.46, CI95%: 0.22-0.82, *Pr*(perturbed (mixed blocks) > non-perturbed (block 1)) = 0.006; OR = 0.38, CI95%: 0.18-0.68, *Pr*(perturbed (mixed blocks) > non-perturbed (mixed blocks) = 0.001))], with no difference between non-perturbed trials of the different blocks (*Pr*(non-perturbed (mixed blocks) > non-perturbed (block 1)) = 0.7), see Fig. 3 a), irrespective of *Session* or *Session Order*. We did not find a meaningful change over time (*Pr*(morning 2 > morning 1) = 0.54) across *Trial Type* and *Session Order*, and no meaningful effect of group (*Pr*(verum-sham ­ sham-verum) = 0.08) across the other factors. We did not find any meaningful interactions, indicating that results did not change from morning 1 to morning 2, neither for the different levels of *Trial Type* nor of *Session Order*.

**Figure 3.**
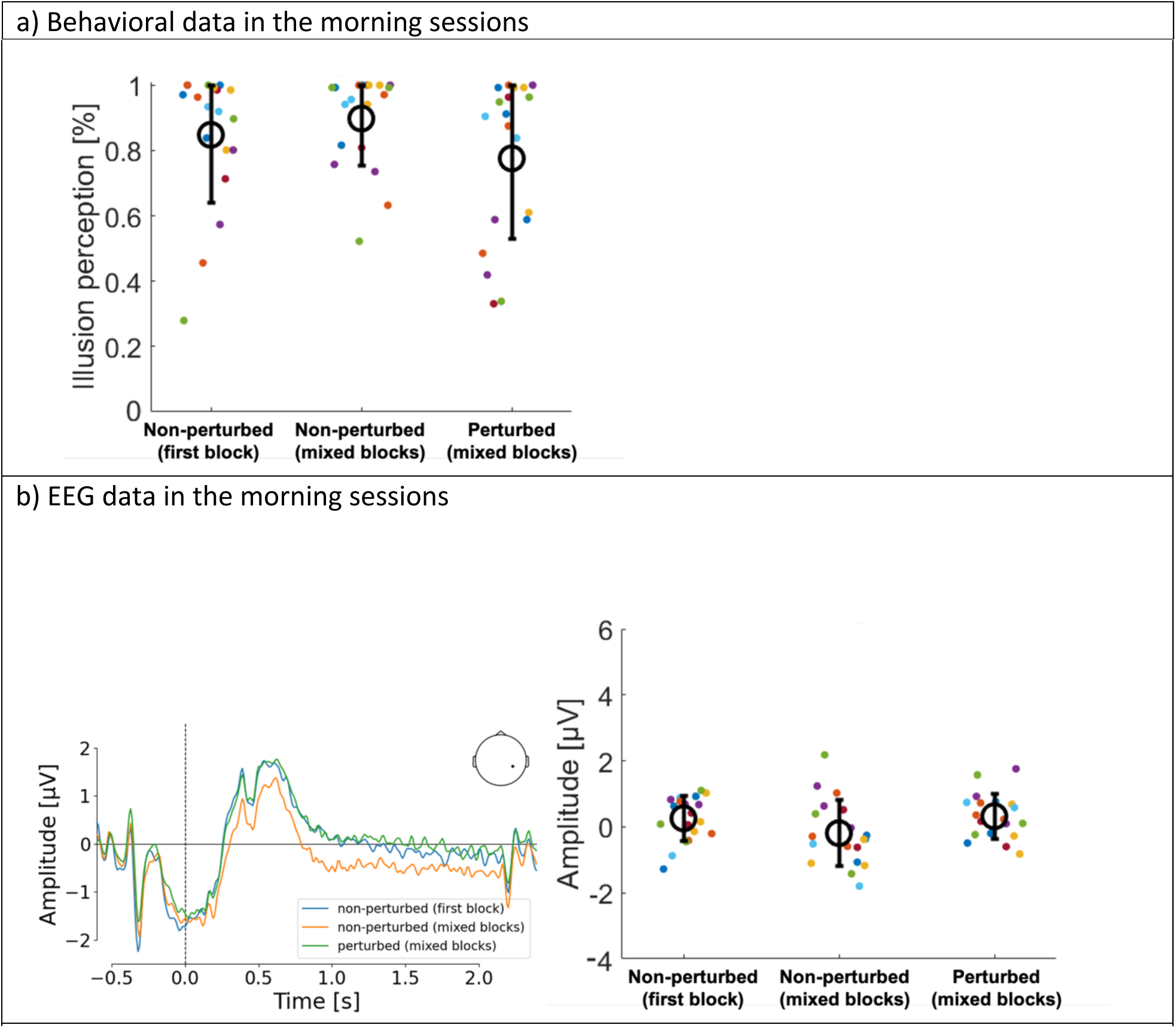
Data from the averaged morning sessions in response to the three trial types. In a) behavioral results are depicted, showing higher illusion perception rates for non-perturbed trials than for perturbed trials. In b) the grand mean ERP traces for electrode CP4 are depicted on the left, showing higher EEG amplitudes for perturbed (green line) than for non-perturbed trials (blue and orange lines). On the right the respective individual EEG peak amplitudes are depicted. In subgraphs depicting individual data, small colored circles represent individual participants, large empty black circles represent the mean and the error bars the standard deviation.

In the EEG data, we found higher ERP amplitudes at electrode CP4 in the time range of 0.15s to 2s after the contact of the squares, for perturbed trials compared to non-perturbed trials of the mixed blocks [OR = 5.5, CI95%: 2.89-9.59, *Pr*(perturbed (mixed blocks) > non-perturbed (mixed blocks) > 0.99)], and compared to non-perturbed trials of the first block [OR = 2.6, CI95%: 1.36-4.6, *Pr*(perturbed (mixed blocks) > non-perturbed (first block)) = 0.99], and smaller amplitudes for non-perturbed trials in the mixed blocks than in the first block [OR = 0.5, CI95%: 0.27-0.83, *Pr*(non-perturbed (mixed blocks) > non-perturbed (first block)) = 0.004], see Fig. 3 b). We did not find a meaningful change over time (*Pr*(morning 2 > morning 1) = 0.48), nor between groups (*Pr*(verum-sham > sham-verum) = 0.78). We found a meaningful interaction between *Trial type* and *Session* (*Pr* = 0.001), which resulted from a decrease in amplitude values from the first to the second morning for perturbed trials (*Pr* < 0.001), which was not there for the non-perturbed trials (*Pr* = 0.48 and *Pr* = 0.965, for non-perturbed trials of the first and the mixed block, respectively). Another meaningful interaction was found between *Trial Type* and *Session Order* (*Pr* = 0.01), together with a meaningful 3-way interaction (*Pr* = 0.99). There were two modulations specific to perturbed trials: 1) in the group sham-verum, amplitudes decreased from the first to the second morning for perturbed trials (Pr < 0.001), which was not the case in the non-perturbed trials (*Pr* = 0.48 and *Pr* = 0.965, for non-perturbed trials of the fist and the mixed block, respectively) and 2) on the second morning, amplitudes where higher for the group verum-sham than for the sham-verum for perturbed trials (*Pr* = 0.98), which was not the case in the non-perturbed trials (*Pr* = 0.78 and *Pr* = 0.47, for non-perturbed trials of the first and the mixed block, respectively).

There was one inter-group difference that did not interact with the session, with smaller amplitudes in non-perturbed trials in the mixed block in the group running TMS in the sham-verum order rather than in the verum-sham order (*Pr* = 0.987). There was no such difference in the first block with non-perturbed trials (*Pr* = 0.78).

The results above confirm an effect of the perturbation on both the illusion rate and the EEG signals. We found meaningful effects suggesting different evolutions across time depending on *Trial types*, which could confound with effects of the TMS interventions. We thus separately investigated the influence of interventions for the three trial types non-perturbed (first block), non-perturbed (mixed blocks), perturbed (mixed blocks). Further, the meaning of a TMS impact on perturbed and non-perturbed trials is different. If TMS changes the illusion itself, e.g., by affecting the processing of contrast prediction errors, then it should affect the results observed in non-perturbed trials. If TMS affects the processing of the trajectory prediction error, then it should affect the results observed in perturbed trials.

### Are responses to the illusion sensitive to TMS interventions?

We investigated the influence of the TMS interventions on the illusion perception rate by means of the factors *Intervention* (verum and sham TMS), and *Time Point* (before and after intervention), separately for the different *Trial Types:* non-perturbed (first block), non-perturbed (mixed blocks), and perturbed (mixed blocks).

The data of the non-perturbed trials in block 1 showed only one marginal effect. The detailed results can be found in Supplementary Material S4.1.

In a next step, we analyzed non-perturbed trials in the mixed blocks and found no effect of *Intervention* (*Pr*(verum TMS > sham TMS) = 0.49), no effect of *Time Point* (*Pr*(after > before) = 0.2), and no meaningful interaction (*Pr* = 0.25). Further, *Session Order* did not show any meaningful effects (*Pr*(verum-sham > sham-verum) = 0.11) nor interactions, see Fig. 4 a). In the EEG data, we did not find a meaningful effect of *Intervention* (*Pr*(verum TMS > sham TMS) = 0.83), but amplitudes increased from before to after TMS [effect of *Time Point*: OR = 2.04, CI95%: 1.28-3.1, *Pr*(after > before) = 0.99] independently of the type of *Intervention.* We additionally found a meaningful interaction between *Intervention* and *Time Point* indicating an increase of amplitudes from before to after sham TMS (*Pr* = 0.99), which was not the case for verum TMS (*Pr* = 0.56), see Fig. 4 b). Regarding *Session Order*, we did not find a main effect (*Pr*(verum-sham > sham-verum) = 0.94), and *Session Order* only interacted with *Time Point* (*Pr* = 0.023), but not with *Intervention* (*Pr* = 0.19), nor was there a meaningful three-way interaction (*Pr* = 0.903).

**Figure 4.**
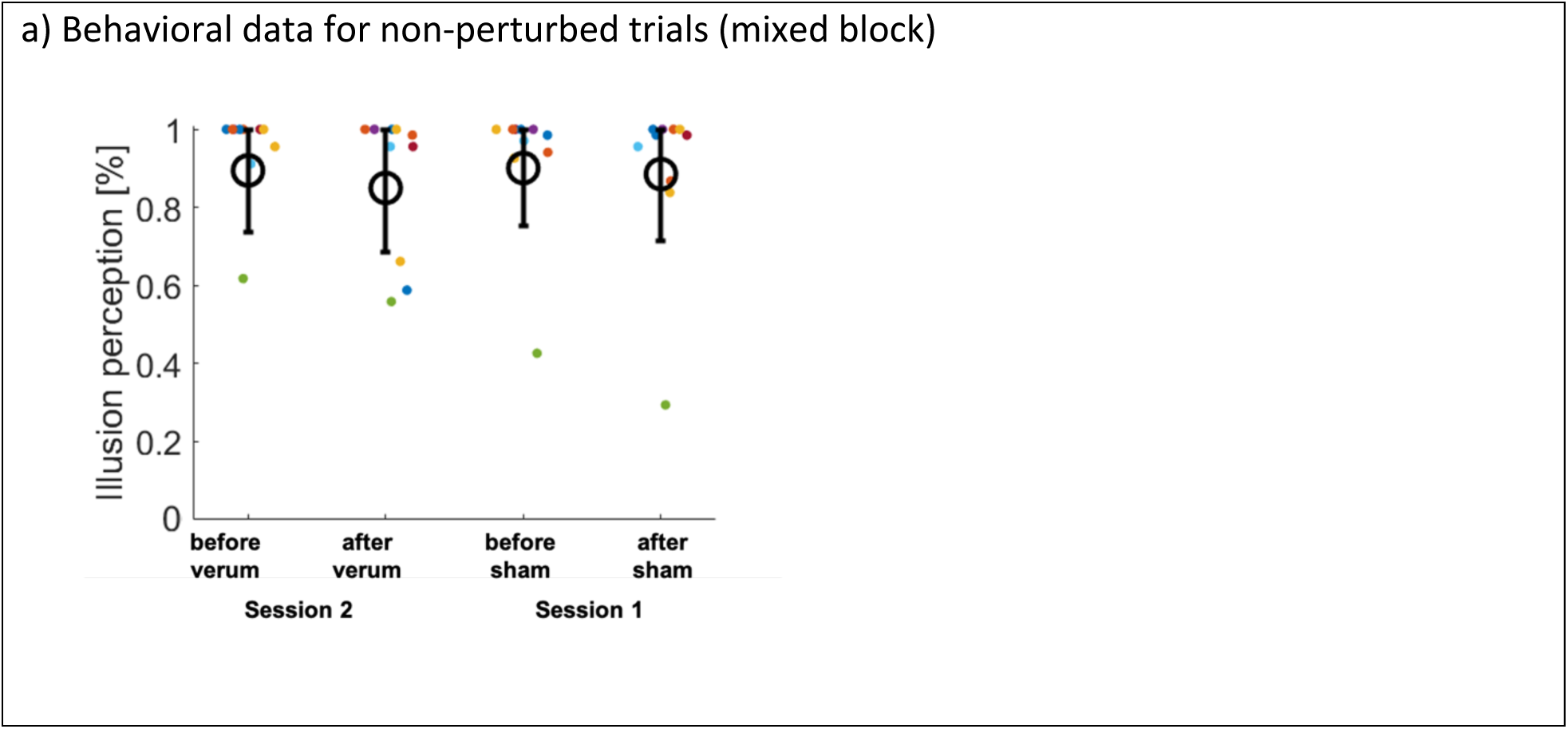
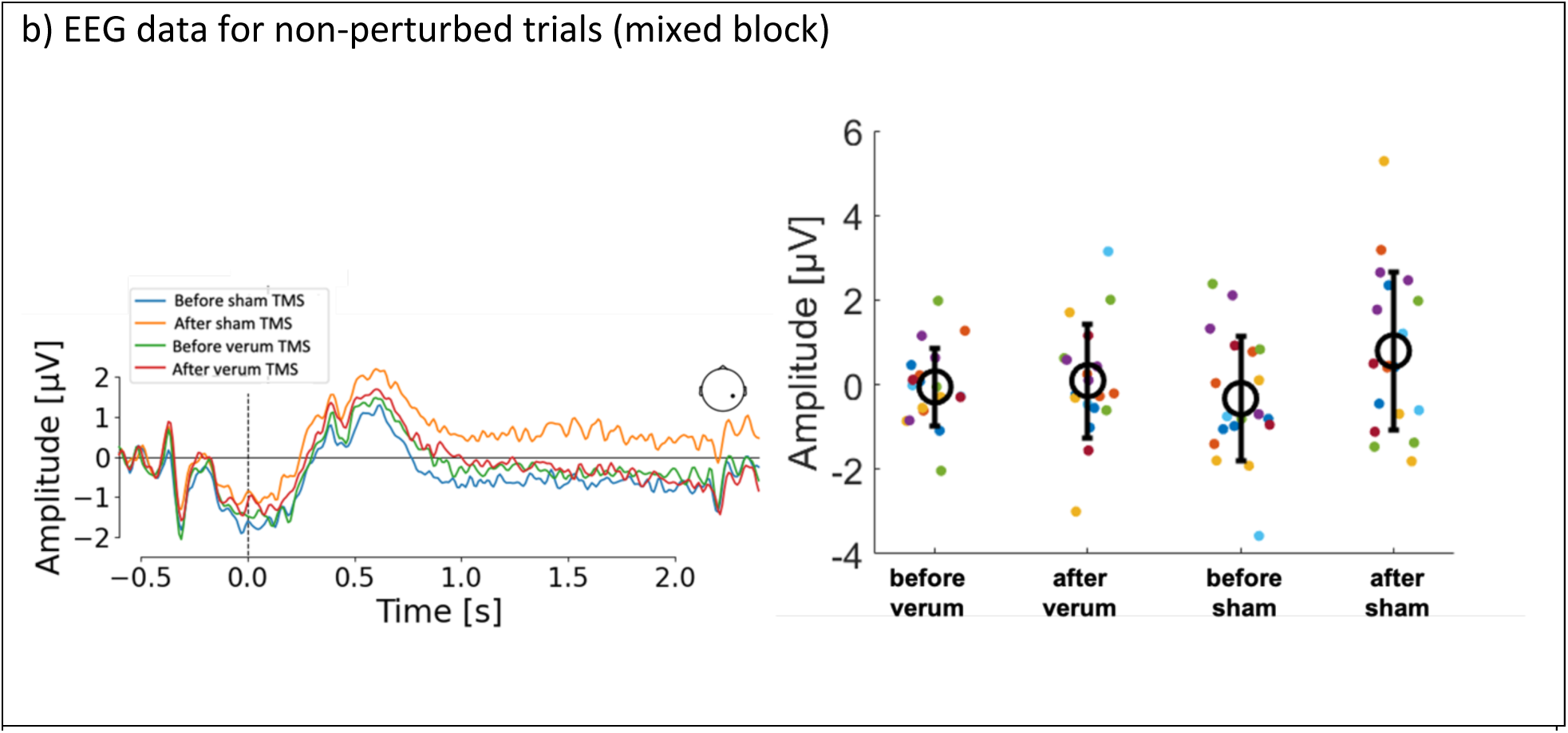
Non-perturbed trials of the mixed blocks – the influence of *Time Point* and type of *Intervention.* In a) behavioral data, the illusion rate was only marginally lower after than before the intervention, but neither the type of *Intervention*, nor an interaction was indicated. In b) EEG data, we depicted grand mean ERP data on the left, while on the right individual is depicted. The meaningful interaction indicated a much stronger increase of amplitudes from before to after sham TMS, compared to verum TMS. Small colored circles represent individual participants’ data, large empty black circles represent the mean and the error bars the standard deviation.

Third, we investigated perturbed trials and found that neither *Intervention* (*Pr*(verum TMS > sham TMS) = 0.26), nor *Time Point* (*Pr*(after > before) = 0.24), nor an interaction (*Pr* = 0.28) had meaningful effects on the illusion rate. *Session Order* did not show a meaningful effect (*Pr* = 0.33), nor meaningful interactions (Pr between 0.06-0.95), see Fig. 5 a). Regarding EEG amplitudes, perturbed trials (see Fig. 5 b) showed several meaningful effects, which can be inspected in Supplementary Material S4.2, but here we want to highlight a clearly meaningful interaction between *Intervention* and *Time Point* (Pr > 0.999), revealing an increase in amplitudes from before to after verum TMS (*Pr*(after verum TMS > before verum TMS) > 0.999), but no such amplitude change for sham TMS (*Pr*(after sham TMS > before sham TMS) = 0.04). Interestingly, we also found a meaningful three-way interaction (*Pr* = 0.012), showing that the increase from before to after verum TMS was found only when it was applied on the second session (*Pr*(after verum TMS for sham-verum > before verum TMS for sham-verum) = 0.999), but not when it was applied on the first session (*Pr*(after verum TMS for verum-sham > before verum TMS for verum-sham) = 0.38), see Fig. 5 b).

**Figure 5.**
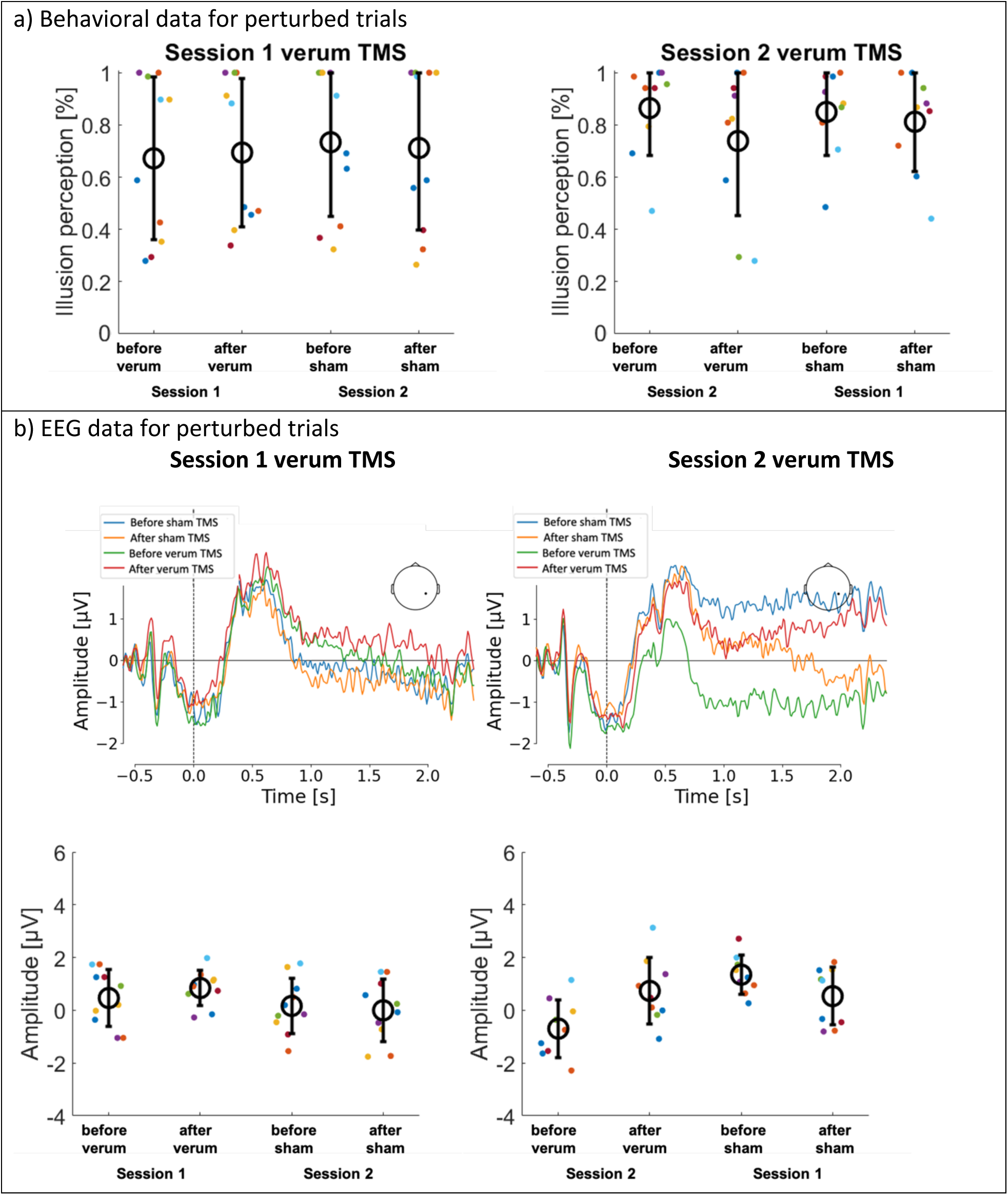
Perturbed trials of the mixed blocks – the influence of *Time Point* and type of *Intervention.* In a) behavioral data, the illusion rate decreased from before to after intervention for the group receiving verum TMS at session 2 (right graph), whereas illusion rates stayed constant for the group receiving verum TMS at session 1 (left graph). In b) EEG data, we depicted grand mean ERP data on the left, while on the right individual is depicted.

We did not find a meaningful three-way interaction in the behavioral results of the perturbed trials, but we did find such an interaction in the EEG, showing clear results for the group sham-verum. In a very exploratory, additional, analysis, we quantified possible behavioral effects for the group sham-verum and found a decrease of illusion from before to after verum TMS intervention (*Pr*(after verum TMS for sham-verum > before verum TMS for sham-verum) = 0.002), but no difference in behavior for the sham TMS intervention (Pr(after sham TMS for sham-verum ­ before sham TMS for sham-verum) = 0.24).

We found for both groups that amplitudes decreased from before to after sham TMS. For verum TMS, amplitudes slightly decreased from before to after in the group receiving verum TMS at session 1 (left graph), but they increased in the group receiving verum TMS at session 2 (right graph). Small colored circles represent individual participants’ data, large empty black circles represent the mean and the error bars the standard deviation.

### Correlation between behavioral and electrophysiological results

We did not find any significant correlations between behavioral and electrophysiological results by means of Spearman rank correlation coefficients in any of the measures.

## Discussion

The main result is that a small and undetectable perturbation of a linear motion trajectory impacts the conscious illusion rate as well as EEG signals. The impact of the perturbation on the illusion rate is stable from the first to the second morning session, while the EEG signals decrease from the first to the second morning session, and specifically so in case of perturbed trials. TMS on the cerebellum modulates EEG responses to perturbed trials, confirming our hypothesis of an involvement of the cerebellum in the effect of the trajectory perturbation. These changes occur depending on the *Session Order* (verum-sham vs. sham-verum). Together with the spontaneous change in EEG signals over time in case of a trajectory perturbation, the results suggest an impact of the stimulation on the modulation of the response to the perturbation rather than an effect on the processing of the perturbation per se.

Subliminal manipulations are known to affect behavioural responses and brain activation, like in priming studies (Pessiglione et al., 2007; Van den Bussche et al., 2009). Previous studies have also shown that sub-threshold prediction errors elicit EEG signals, often early ones (Rowe et al., 2020; Teixeira et al., 2020). It is usually believed that the processing of sub-threshold prediction errors can help adjusting learning (to perceive or to move) (Bavassi et al., 2013; Rose et al., 2005). To which amount such errors have an impact on consciousness is less clear (Foerster et al., 2021; Weibel et al., 2013). In itself the illusion used in the present study shows the conscious consequence of a sub-threshold prediction error (Jovanovic et al., 2023). As a reminder, we hypothesized that the regularity of the trajectory allows for the illusion, by making it possible to predict the future position of the two squares, together with the figure-ground contrast of their leading edge. When the two squares touch, the figure-ground contrast disappears, as the two squares are drawn with the same gray level (see Jovanovic et al., 2023 (Jovanovic et al., 2023) for different, control, situations). The disappearance of the figure-ground contrast represents a prediction error, but this error is not consciously perceived. On the contrary, participants do not see the contact, and instead see a gap between the squares.

The fact that a perturbation of the trajectory reduces the illusion supports the hypothesis that the trajectory regularity plays a crucial role in the illusion. What is noteworthy in the present results is that the impact of the trajectory manipulation is almost immediate. When the square jumps by 2 instead of 1 pixel, it takes 17ms, and 17 ms for a last 1-pixel-move. The square then stops for 30 ms and disappears. There is thus 68 ms between the start of the jump and the disappearance of the square. Despite this short delay, the jump appears to facilitate the detection of a contact between the two squares. It is unlikely an attention effect. First, the jump is undetectable (see Supplementary Material S3) and 68 ms is too short to displace attention. It is also too short to allow for a top-down modulation effect (Supèr et al., 2001). Additionally, the jump means that the squares disappear 17 ms earlier than in the non-perturbed trials. This should have made it more difficult to see the contact between the two squares. It is thus all the more remarkable that the illusion decreases, meaning that the participants see the contact more frequently in case of a perturbed trajectory. Although counter-intuitive, the results are consistent with the idea that the trajectory perturbation disturbs the trajectory prediction itself. The 68 ms delay before squares’ disappearance does not allow to readjust the prediction online, but the disturbance of the trajectory prediction would make it more difficult to emit a prediction error and thus to evoke the illusion.

The decrease in the illusion rate was accompanied by an increase in a positive EEG signal, which looks like a late positive potential (LPP) (Hajcak & Foti, 2020), at least the type of LPP signal that is observed in case of a neutral stimulus, i.e., with a slowly decreasing amplitude. The observation of a LPP may appear as surprising at first sight. The LPP is mainly described in response to emotional prediction error, i.e., threat (Botelho et al., 2023; Del Popolo Cristaldi et al., 2021). The amplitude of the LPP is increased by motivation and emotion, which does not seem to play a major role in our study. Nonetheless, some studies show a LPP independently from an emotional response (Hajcak et al., 2010; Hofmann et al., 2019; Jahshan et al., 2015; MacKay et al., 2024; Sun et al., 2017). Interestingly, two studies used paradigms with moving stimuli, MacKay et al. (2024 (MacKay et al., 2024)) used implied movement, and Jahshan et al. (2015 (Jahshan et al., 2015)) used biological motion. However, to the best of our knowledge, no study recorded a LPP related to a trajectory manipulation, and at this stage it is hard to know if motion plays a significant role in the LPP effect. What may be especially pertinent is Bradley’s proposal (2009 (Bradley, 2009)) that the LPP amplitude is an indicator of stimulus significance, which can be due to novelty, emotion or task relevance (Hajcak & Foti, 2020). In the literature, the LPP is often elicited with rare targets, like in oddball tasks resulting in a mismatch negativity. In contrast, the perturbed trials in our task represent half of the trials. Nonetheless, as already pointed out, a change in trajectory might be critical, e.g., on the road, and a prediction error due to a change in trajectory should represent a salient information. One of the consequences of predictive coding is precisely to increase the saliency of pertinent events (Chalk et al., 2018; Kanai et al., 2015). A first possible explanation is that in our study the saliency does not make the perturbation conscious, but might have helped to adjust how the following trajectory is predicted, e.g., by (not-consciously) integrating the slight acceleration in the prediction. If the trajectory prediction is correct, i.e., when the following trajectory integrates a slight acceleration, there is no prediction error and the illusion can arise as in non-perturbed trials. This adjustment should thus have led to significant trial-to-trial effects in behavioural responses, which was not the case, see Supplementary Material S4.3. The randomization of the trials probably makes this type of adjustment inefficient. The adjustment would be useful only in half cases after a perturbation, and would require an adjustment in the opposite direction after a non-perturbed trial. The LPP unlikely reflects prediction adjustment directly. According to the literature, the LPP may play an alerting role in signaling the pertinence of a signal, in our case the trajectory perturbation. The pertinence of the signal may be required for an adjustment to take place, but if this adjustment is not really efficient, the saliency of the perturbation may decrease over time. This might explain the decrease in the EEG signal from the first to the second morning in case of a perturbation.

In all, the results in the morning sessions before TMS take place, suggest that the perturbation is not ignored, and is accompanied by an EEG signal conveying its significance. The perturbation likely impairs the trajectory-related prediction, which may explain the drop in the illusion rate, as the contact between squares occur a very short time after the adjustment, when the prediction could not have been reinstated. In fact, it suggests that for a very short time there might be a lack of prediction. The perturbation still has a behavioural effect on session 2, showing that it is difficult to ignore, probably because perturbed and unperturbed trials are randomized. Its significance may vane with time, however, as suggested by the decrease in the LPP in the second morning session relative to the first one. This decrease might play a role in the impact of TMS.

It should be emphasized that the mere existence of an impact of TMS in this precise study rather came as a surprise. Given the potential side effects of a stimulation of cerebellum in humans, we had been very cautious by using a short and single stimulation. As a matter of fact, stimulating the cerebellum is reputed to be difficult due to its proximity to the neck muscles and to the vestibular system, potentially inducing muscle contractures, dizziness or nausea (Hurtado-Puerto et al., 2020; Satow et al., 2002), although Hurtado-Puerto et al. (2020 (Hurtado-Puerto et al., 2020)) showed that theta-burst stimulations were well tolerated, with around 4% of side-effects occurrences. It should be emphasized that in the present study no side-effect was observed, probably thanks to careful neuro-navigation and the use of a TMS-robot. Most importantly, and despite the limited duration of stimulation we observed an impact of TMS, which was discrete on behaviour, but clearer on EEG effects.

It should be first noted that in the absence of any perturbation, most effects of the TMS on the illusion rate or EEG signals were marginal and differed in the first block and the mixed blocks. We did not expect that TMS on the cerebellum would affect the illusion itself, given figure-ground contrast is processed elsewhere than in the cerebellum. In fact, the only clearly significant results of TMS were found on EEG signals in the mixed block, and appear to mirror the effects in case of a trajectory perturbation. In case of a perturbation, the LPP amplitude significantly increased after the stimulation, at least in participants being stimulated with a verum TMS on session 2. When conducting a very exploratory analysis in this group (sham-verum), we found a decrease of illusion rate after the stimulation. In contrast, in case of non-perturbed trials in the mixed blocks, the LPP decreased after the sham TMS, but remained stable after the verum TMS. The association between an increase in LPP and drop in the illusion rate for perturbed trials match the results observed in session 1 in the morning. This parallelism suggests that the stimulation of the cerebellum led to increased responsiveness to the millisecond-level trajectory perturbation. Such an impact coheres with the literature suggesting a role of the cerebellum in the processing of prediction errors (Brooks & Cullen, 2013), motion (Ivry* & Diener, 1991; Thier et al., 1999), motion trajectory prediction (Baumann et al., 2015; Baumann & Mattingley, 2010; O’Reilly et al., 2008), and millisecond-level timing (Coull et al., 2011; Ivry & Spencer, 2004). Given the known roles of the cerebellum, the results further reinforce our interpretation that the illusion comes about as a result of trajectory prediction, and decreases due to the millisecond-level trajectory perturbation.

Despite the coherence of the results with the literature, and the use of one single and short-duration stimulation, which likely reduced its impact, the effect of *Session Order* should be discussed. As a matter of fact, no clear impact of the TMS is observed in those participants who had a verum TMS on the first session. A general group effect is not supported by the data since the group did not reveal meaningful effects in any of the behavioral analyses, except for the special case of perturbed trials described above. An explanation may thus be found in relation with the LPP. As we have seen the LPP appears to reflect the significance of the perturbation. If we follow this reasoning, TMS would result in an increase of the saliency of the perturbation. However, this increase was mainly examined relative to the LPP amplitude recorded in the second morning session, which had decreased relative to the first session. If the LPP decrease was due to a decrease in the significance of the perturbation, as suggested above, then TMS may have mainly reinstated a significance that had worn off.

The fact that a cerebellum stimulation has an impact on a saliency effect is not as surprising as it might seem. First the role of cerebellum in emotion processing is now recognized (Schmahmann, 2010), and the cerebellum has also been suggested to play a role in saliency (Nguyen et al., 2017), possibly through interactions with amygdala (Taub & Mintz, 2010). Cerebellar lesions have been suggested to induce a decrease in the LPP evoked by emotional pictures (Adamaszek et al., 2015). Moreover, as already emphasized in the introduction, the cerebellar cortex, in particular Purkinje cells discharges and climbing fibers control have been proposed to play a role in the encoding of prediction errors (Kimpo et al., 2014; Streng et al., 2022). The modulation of firing rates in the cerebellar cortex triggered by TMS illustrates how processing of prediction errors can be modulated.

In conclusion, we provide evidence that the manipulation of a square’s moving trajectory at the millisecond-level affects the processing of the trajectory within the 100 ms following the trajectory, very probably by interrupting the trajectory prediction. This effect occurs even as the manipulation remains sub-threshold. Despite the sub-threshold and non-conscious character of the manipulation, it affects conscious perception, as evidenced by the consequence on the rate of the illusion. The manipulation further impacts EEG signals, in the form of an LPP. Moreover, the stimulation targeting the CRUS I/II in the cerebellum affects behaviour slightly and more clearly the LPP, at least in those participants who received verum TMS after having performed the task three times. As LPP decreases over time, the results suggest the effect of TMS may represent a reinstation of the EEG consequences of the prediction error, i.e., a modulation of its significance. Whether this is a beneficial effect or not remains to be explored. As a matter of fact, the randomization of the perturbed and non-perturbed trials made it difficult to adjust to the perturbation from trial to trial, and thus to learn from the perturbation. It is all the more remarkable that the behavioural impact of the trajectory manipulation did not fade like the LPP did. The results suggest that the acceleration cannot be ignored. It might be impossible not to process the perturbation at least initially, and this would be visible in our task because the impact of the perturbation occurs in less than 100 ms after the acceleration. The LPP, on the other hand occurs much later, and may reflect the suppression of the signal, when it decreases over time.

## Acknowledgements

This work was funded by an Agence National de Recherche grant (ANR-16-CE37-0004-02) awarded to A.G, a Fondation de France grant (RAF17001 MMA) awarded to A.G and P.I., a Strasbourg University Idex ‘Projet Exploratoire 2019’ grant (EOTP W19RHUS4) awarded to A.G, P.I. and J.F., and a Neurotechnology Inserm Boost grant (R24057MM) awarded to A.G.. E.J. was financed by the EU, Horizon 2020 Framework Program, FET Proactive (VIRTUALTIMES consortium, grant agreement Id: 824128 to Anne Giersch), the Deutsche Forschungsgemeinschaft grant (KO 4764/9-1, TE 280/26-1), and the Institute for Frontier Areas of Psychology and Mental Health, Freiburg, Germany. The authors would like to thank Corinne Marrer and Romane Weill-Rossi for the work in the MRI sessions. Further, we greatly appreciate the work of the employees at the side for TMS (CEMNIS), in particular Mandy Haumesser, Mathieu Ricka, Tiffany Schilt, and Ludovic Dormegny-Jeanjean. At last, we want to thank Estelle Koning, who took part in the complex organization of the study.

