## Supplementary Material S1, S2, S3, S4 for "Subthreshold violations of trajectory predictions are sensitive to TMS of Cerebellum CRUS I/II"

### Supplementary Material for: Subthreshold violations of trajectory predictions are sensitive to TMS of Cerebellum CRUS I/II

#### Supplementary Material S1: Does the Target location influence consequences of TMS of the Cerebellum?

The cerebellum is a deep structure that is difficult to stimulate using transcranial magnetic stimulation (TMS). In detail, we wanted to stimulate the border between Crus I and Crus II. Due to the differences in individual scalp properties, we preselected two possible targets. Target 1 was the most lateral and more anterior part of the border between Crus I and II and was targeted at first. In case target 1 was not accessible in the individual, we stimulated target 2, a more lateral target that is closer to the intermediate regions. We successfully stimulated target 1 in 19 participants and target 2 in 5 participants. Here we aim at verifying that the target location did not have any systematic influence on the results.

As described in the main manuscript, we excluded three participants due to abnormal responses (see Behavioral analysis of the methods section in the main manuscript). Two of those participants were stimulated at target 1 and one was stimulated at target 2. Therefore, were able to compare results between target 1 and target 2 groups from 17 and 4 participants, respectively. We did not find any differences using Wilcoxon signed rank tests between the two groups for the behavioral results (illusion perception rate) before the two types of interventions for neither non-perturbed trials, nor for perturbed trials. From this, we concluded that target location had no systematic influence on the results of interest and did not regard this factor in the analyses presented in the main manuscript.

#### Supplementary Material S2: Trial type selection

In the collision task, two squares move towards each other until their inner edges touch in the middle of the screen. In a previous study(Jovanovic et al., 2023), it was found that in case of short (17ms and 33ms) contact durations this can result in an illusory perception of a gap between the two squares.

We did not have any a priori assumptions to decide whether the TMS intervention would result in an increase or in a decrease of the illusion perception rate in case of a trajectory manipulation. In a previous study(Jovanovic et al., 2023) (see also Fig. S1), the illusion perception rate for 17ms contact duration was 89.6% (SD: 3.9%) and for 33ms contact duration it was 67.1% (SD: 7.5%). We thus selected the 33ms contact duration trials to constitute the majority of trials in the currently conducted study, allowing us to reliably measure both a possible increase or decrease of the illusion perception rate.

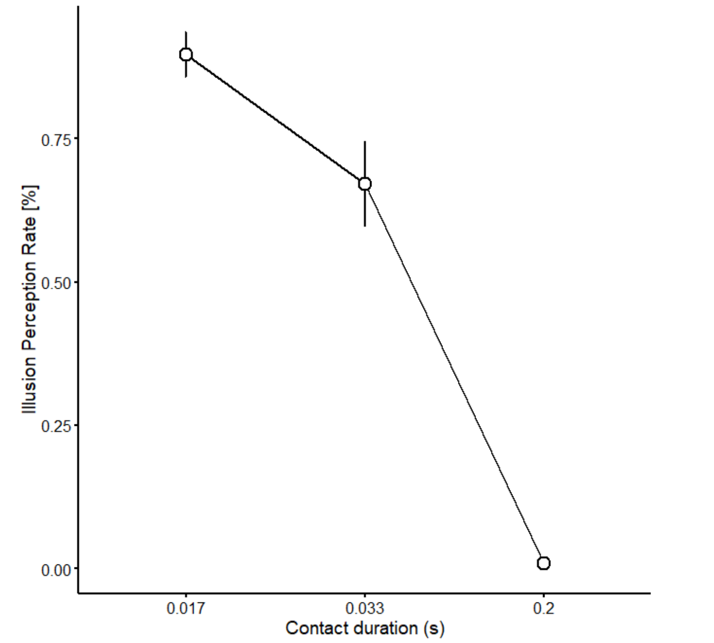


Fig. S1 Illusion in a previous study. The illusion perception rate is decreasing with increasing contact duration. Circles represent the mean value across 12 participants and error bars the standard deviation.

#### Supplementary material S3: Perturbation detection

Perturbations of the squares' trajectory take place 2 frames before they contact each other in the middle of the screen. In a separate control experiment, we asked participants to perform, after each stimulus, the illusion task ("did the squares touch?") and in a second task to indicate whether they perceived a perturbation or not. With this we aimed to test whether the perturbation reached a conscious level. We presented the same ratio of 17ms, 33ms, and 200ms trials as in the perturbation condition described in the main manuscript, but we only presented 100 trials overall, with half of the trials containing a perturbation and the other half not. In order to not bias participants towards consciously perceiving perturbations in the main results, we performed this control experiment at the end of the second measurement day, i.e. after finishing all the other experimental conditions.

As can be seen in Fig. S2, participants do not show different rates of perturbation perception for trials containing perturbations or not (Wilcoxon signed rank test: $Z$=-1.14, $p$=0.26), suggesting that they did not reliably detect perturbations. Given these results, it is justified to compare results of perturbed and non-perturbed trials, as reported in the main manuscript.

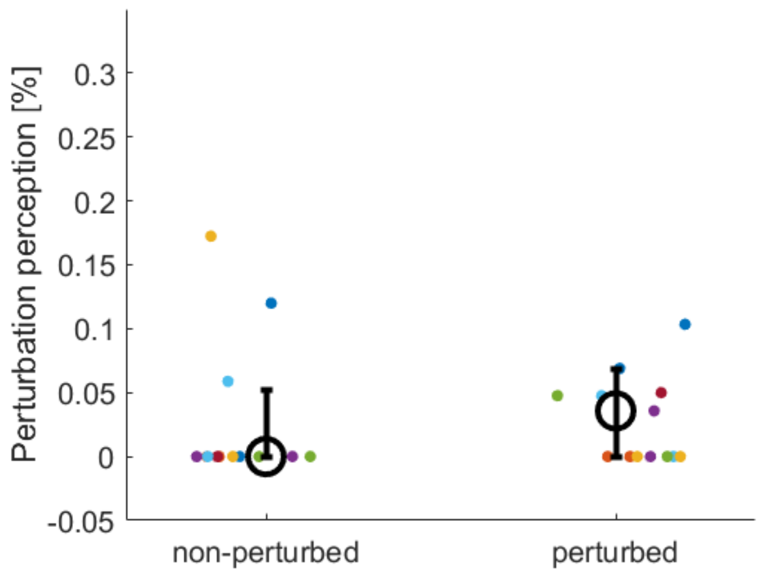


Fig. S2 The rate of perturbation perception in a control condition is displayed for 33ms trials. On the left the control trials with no perturbation show the baseline for perturbation perception in case of its absence. On the right, the trials including a perturbation are presented. Small colored circles represent individual data, large black empty circles the median and the error bars the interquartile ranges.

#### Supplementary Material S4: Influences of TMS interventions on responses to the illusion

##### *S4.1 Non-perturbed trials of the first block*

Here, we considered only non-perturbed trials of the first block, see Fig. S3 a). We found no influence of *Intervention* (*Pr*(verum TMS > sham TMS) = 0.34), and no effect of time (*Pr*(after > before) = 0.47). Further, we found no interaction effect (*Pr* = 0.15). Regarding the two groups, we found no effect of *Session Order* (*Pr*(verum-sham > sham-verum) = 0.09), and no meaningful interactions, despite the graph suggesting otherwise (*Pr*s between 0.3-0.85). The EEG results showed no effect of *Intervention* (*Pr*(verum TMS > sham TMS) = 0.72) and no effect for *Time Point* (*Pr*(after > before) = 0.12). We found a marginal interaction effect (*Pr* = 0.952), indicating no difference between verum and sham TMS before the intervention (Pr(verum TMS before > sham TMS before) = 0.72), but larger amplitudes for verum TMS than for sham TMS after the intervention (Pr(verum TMS after > sham TMS after) = 0.99), see Fig. S3 b). When considering *Session Order*, we did not find any clearly meaningful effects nor interactions.

a) Behavioral data for non-perturbed trials (block 1)

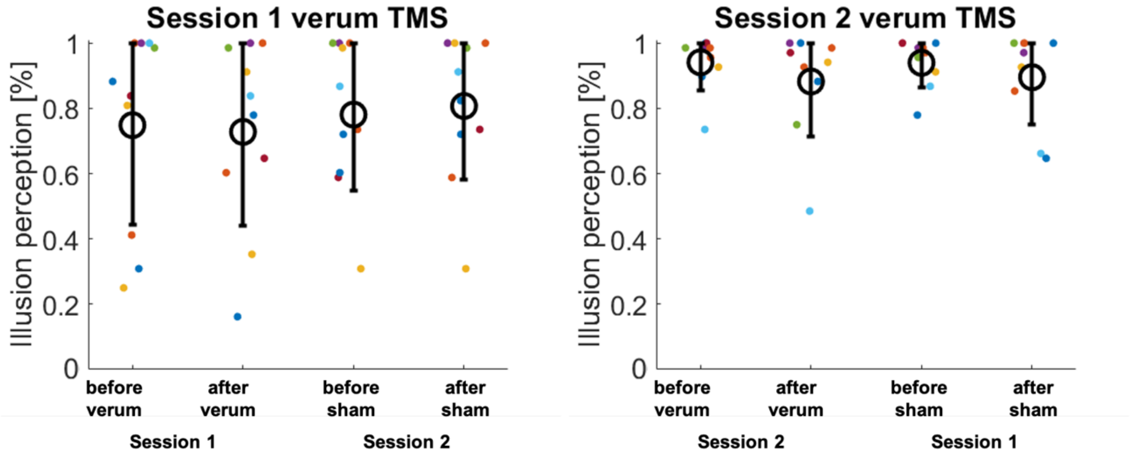


b) EEG data for non-perturbed trials (block 1)

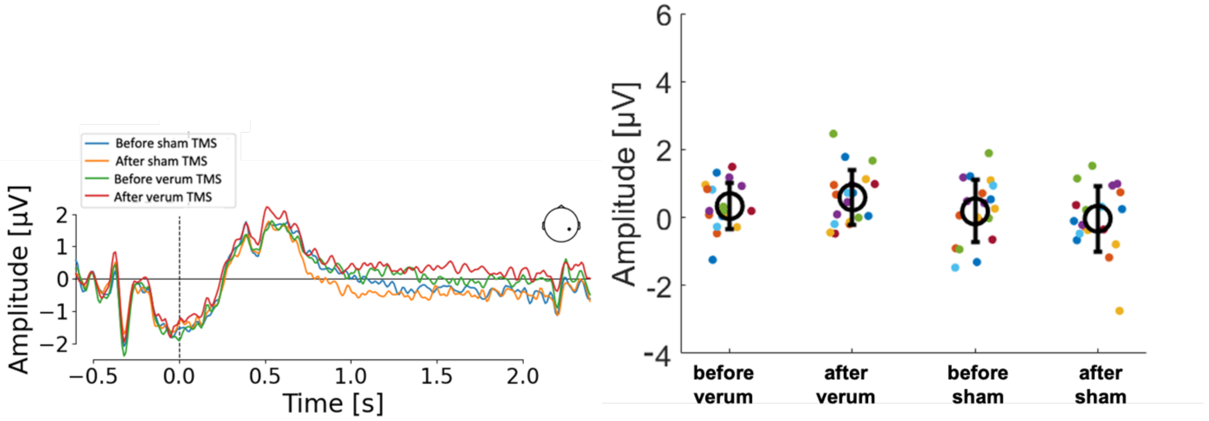


Figure S3 – Non-perturbed trials of the first block – the influence of *Time Point* and type of *Intervention.* In a) behavioral data, the illusion rate marginally decreased from before to after verum TMS, but stayed the same for sham TMS. On the left, the group receiving verum TMS on session 1 is depicted, which shows marginally smaller illusion rates than the group receiving verum TMS on session 2, depicted on the right. In b) EEG data, we depicted grand mean ERP data on the left, while on the right individual is depicted. A marginal interaction indicated an amplitude increase from before to after verum TMS, while there was a slight amplitude decrease for sham TMS. Small colored circles represent individual participants' data, large empty black circles represent the mean and the error bars the standard deviation.

#### *S4.2 Perturbed trials – EEG results*

Investigating EEG amplitudes in response to the perturbed trials, we found clearly higher amplitudes for sham TMS than for verum TMS [OR = 0.19, CI95%: 0.09-0.37, *Pr*(verum TMS > sham TMS) < 0.001], when averaged over before and after intervention. Furthermore, marginally higher amplitudes before than after intervention (*Pr*(after > before) = 0.04), and higher amplitudes for the group sham-verum than for verum-sham [OR = 0.44, CI95%: 0.18-0.89, *Pr*(verum-sham > sham-verum) = 0.012] were found when averaged across the respective other factors. As described in the main manuscript, we found a clearly meaningful interaction between *Intervention* and *Time Point* (*Pr* > 0.999), which was due to the following two meaningful effects in the pairwise comparisons: 1) amplitudes increased from before to after verum TMS (*Pr*(after verum TMS > before verum TMS) > 0.999) and 2) amplitudes were higher before sham TMS than before verum TMS (*Pr*(before verum TMS > before sham TMS) < 0.001), while no other comparisons were meaningful. Further, we found a clearly meaningful interaction effect between *Intervention* and *Session Order* (*Pr* > 0.99), which was due to three meaningful effects in the pairwise comparisons: 1) in the group that received verum TMS on the second day (group sham-verum), amplitudes decreased from before to after intervention (*Pr*(after for sham-verum > before for sham-verum) < 0.001), 2) we found larger amplitudes before the intervention for group sham-verum than for verum-sham (*Pr*(before verum-sham > before sham-verum) = 0.012), but 3) we found larger amplitudes after the intervention for group verum-sham than for sham-verum (*Pr*(after verum-sham > after sham-verum) = 0.99). We did not find an interaction between *Time Point* and *Session Order* (*Pr* = 0.82), but as described in the main manuscript, we found a clearly meaningful three-way interaction (*Pr* = 0.012). Pairwise comparisons for this interaction were traced back to 13 meaningful pairwise comparisons that are listed below.

Meaningful comparisons for the group sham-verum:

- *Pr*(after verum TMS for sham-verum > before verum TMS for sham-verum) = 0.99
- *Pr*(before verum TMS for sham-verum > before sham TMS for sham verum) < 0.001

Meaningful comparisons for the group verum-sham:

- *Pr*(before verum TMS for verum-sham > before sham TMS for verum-sham) < 0.001
- *Pr*(after verum TMS for verum-sham > after sham TMS for verum-sham) = 0.004
- *Pr*(after verum TMS for verum-sham > before sham TMS for verum-sham) = 0.004

Meaningful comparisons between groups:

- *Pr*(before sham TMS for verum-sham > before sham TMS for sham-verum) = 0.012
- *Pr*(before verum TMS for verum-sham > before verum TMS for sham-verum) = 0.012
- *Pr*(after sham TMS for verum-sham > after sham TMS for sham-verum) = 0.012
- *Pr*(after verum TMS for verum-sham > after verum TMS for sham-verum) = 0.005
- *Pr*(before verum TMS for verum-sham > before sham TMS for sham-verum) < 0.001
  *Pr*(after verum TMS for verum-sham > after sham TMS for sham-verum) = 0.005
- *Pr*(after verum TMS for verum-sham > before sham TMS for sham-verum) = 0.005
- *Pr*(after sham TMS for verum-sham > before sham TMS for sham-verum) = 0.009

*S4.3 Trial-to-trial effects:*

It is widely known that previous trials influence the perception of a current trial. In this analysis step, we investigated the influence of the perturbation level of a previous trial N-1 on the illusion perception rate of a current trial N, separately for its stimulus type (non-perturbed, perturbed). Analyses were conducted the same way as described in the main manuscript, i.e., with the factors *Intervention* (verum and sham TMS), *Time Point* (before and after intervention), and *Session Order* (group sham-verum, group verum-sham), and with the additional factor previous *trial N-1* (no perturbation or perturbation, averaged over the previous trials' contact durations).

We did not find any clearly meaningful influence nor interaction of the identity of trial N-1 for the behavioral data (all *Pr* ranging between 0.11– 0.86), neither for the non-perturbed trials nor the perturbed trials at timepoint N. Regarding EEG data, we did not find any meaningful effects nor interactions of the identity of trial N-1 for the non-perturbed trials at timepoint N (all *Pr* ranging between 0.16-0.85). When analyzing the EEG data of perturbed trials at timepoint N, we found no meaningful main effect, nor any interaction including *Intervention* and *Time Point*. We did find some interactions with the other factors, detailed below, which, however, do not contribute to understand the meaningful 3-way interaction for EEG data related to perturbed trials, as described in the last result chapter of the main manuscript (see also Fig. 5 b).

Investigating EEG amplitudes in response to perturbed trials at timepoint N in an analysis considering also the identity of trial N-1, we did not find any main effect of *trial N-1 (Pr* = 0.97), nor the critical interaction between the factors *trial N-1*, *Intervention*, and *Time Point* (*Pr* = 0.13), nor a meaningful 4-way interaction including *Session Order* (*Pr* = 0.87). We did, however, find a meaningful interaction between *trial N-1* and *Time Point* (Pr = 0.998), where amplitudes decreased from before to after intervention when trial N-1 was non-perturbed (*Pr* = 0.000073), but no amplitude change when trial N-1 was perturbed (*Pr* = 0.63). We found one more meaningful interaction including *trial N-1*, which was with the factors *Time Point* and *Session Order* (*Pr* = 0.003). Pairwise comparisons revealed an amplitude decrease from before to after intervention for non-perturbed trials at N-1 in the group sham-verum (*Pr* = 0.000073), which was not the case for perturbed trials at N-1 in the same group (*Pr* = 0.63). In the group verum-sham, we found this decrease of amplitudes from before to after intervention for both, the non-perturbed trials at N-1 (*Pr* = 0.000073), as well as for the perturbed trials at N-1 (*Pr* = 0.0018). In addition to these 4 pairwise comparisons, we found the following meaningful results when understanding the 3-way interaction:

- Pr(before non-perturbed for sham-verum > before non-perturbed for verum-sham) = 0.014
- Pr(before perturbed for sham-verum > before perturbed for verum-sham) = 0.014
- Pr(after non-perturbed for sham-verum > after non-perturbed for verum-sham) = 0.014
- Pr(after perturbed for sham-verum > after perturbed for verum-sham) = 0.002
- Pr(before non-perturbed for verum sham > after perturbed for sham-verum) = 0.998
- Pr(before non-perturbed for sham-verum > before perturbed for verum-sham) = 0.023
- Pr(before non-perturbed for sham-verum > after non-perturbed for verum-sham) = 0.0003
- Pr(before non-perturbed for sham-verum > after perturbed for verum-sham) = 0.0025
- Pr(before perturbed for sham-verum > after perturbed for verum-sham) = 0.0014
- Pr(after non-perturbed for sham-verum > after perturbed for verum-sham) = 0.026
- Pr(before non-perturbed for verum-sham > after perturbed for verum-sham) = 0.003
